# Heterozygous mutation of Sonic Hedgehog receptor (Ptch) drives cerebellar overgrowth and sex-specifically alters hippocampal and cortical layer structure, activity, and social behavior in female mice

**DOI:** 10.1101/2020.01.25.919506

**Authors:** Thomas W. Jackson, Gabriel A. Bendfeldt, Kelby A. Beam, Kylie D. Rock, Scott M. Belcher

**Affiliations:** Center for Human Health and the Environment, Department of Biological Sciences, North Carolina State University, 127 David Clark Labs Campus Box 7617, Raleigh, North Carolina, USA 27695-7617

**Keywords:** cerebellum, cortex, holoprosencephaly, hyperactivity, medial prefrontal cortex, medulloblastoma, sex differences

## Abstract

Sonic hedgehog (SHH) signaling is essential for the differentiation and migration of early stem cell populations during cerebellar development. Dysregulation of SHH-signaling can result in cerebellar overgrowth and the formation of the brain tumor medulloblastoma. Treatment for medulloblastoma is extremely aggressive and patients suffer life-long side effects including behavioral deficits. Considering that other behavioral disorders including autism spectrum disorders, holoprosencephaly, and basal cell nevus syndrome are known to present with cerebellar abnormalities, it is proposed that some behavioral abnormalities could be inherent to the medulloblastoma sequalae rather than treatment. Using a haploinsufficient SHH receptor knockout mouse model (*Ptch1^+/-^*), a partner preference task was used to explore activity, social behavior and neuroanatomical changes resulting from dysregulated SHH signaling. Compared to wild-type, *Ptch1^+/-^* females displayed increased activity by traveling a greater distance in both open-field and partner preference tasks. Social behavior was also sex-specifically modified in *Ptch1^+/-^* females that interacted more with both novel and familiar animals in the partner preference task compared to same-sex wild-type controls. Haploinsufficency of PTCH resulted in cerebellar overgrowth in lobules IV/V and IX of both sexes, and female-specific decreases in hippocampal size and isocortical layer thickness. Taken together, neuroanatomical changes related to deficient SHH signaling may alter social behavior.

## 1. Introduction

The cerebellum is a brain region important for coordinating control of voluntary motor movement, muscle tone, and balance (Altman and Bayer, 1997). Additionally, the cerebellum is involved in higher-order cognitive functions and related behaviors (Rogers et al., 2013a). Developmental cerebellar damage and abnormal cerebellar structure can result in impairment of motor function, cognition, and social reward behavior. Abnormalities of the prefrontal cortex, thalamus, and some cerebellar structures are a commonly observed feature of autism spectrum disorder (ASD). Emerging evidence supports the existence of behavioral circuits integrating the cerebellum, prefrontal cortex, and thalamus through dopaminergic signaling (Rogers et al., 2013a).

Normal cerebellar function depends on properly integrated actions of neurons residing in the three distinctive layers of the mature cerebellum: the molecular layer (ML), Purkinje cell layer (PL), and the internal granular layer (IGL). The ML is primarily comprised of synaptic interfaces between dendritic arbors of Purkinje cell neurons and the axonal parallel fibers of mature granule cells in the IGL. The PL is demarcated by a monolayer of Purkinje cell bodies that divide the ML from the IGL (Altman and Bayer, 1997). Granule cell precursors (GCPs) arise from a population of rhombic lip progenitors that migrate to the cerebellum and form the external germinal layer (EGL) where GCP proliferation continues until post-natal day 15 in mouse (Altman and Bayer, 1997). During the postnatal period of cerebellar development, the GCPs mitotically arrest, differentiate, and migrate through the ML and PL, and then reach their final destination in the IGL as mature granule cells. Development of the stereotypic structure of the cerebellum is a tightly regulated process that requires specific gradients of key morphogens (Martinez et al., 2013).

The Sonic Hedgehog (SHH) signaling pathway is necessary to induce differentiation and mitotic arrest of GCPs in the EGL, and subsequently acts to direct migration of the maturing granule cells to the IGL (Dahmane and Ruiz i Altaba, 1999). Those processes are mediated by the morphogen SHH which binds to the Patched-1 (PTCH1) receptor expressed in Purkinje cells, GCPs, and cerebellar interneurons. Secretion of SHH from Purkinje cells is initiated around E17.5 in the mouse and peaks around postnatal day 6-8. The resulting SHH concentration gradient regulates differentiation of GCPs wherein lower concentrations of the morphogen allow continued proliferation. In the absence of the SHH ligand, PTCH1 inhibits Smoothened (SMO), a downstream G-protein coupled receptor. By contrast, SHH binding leads to disinhibition of SMO, resulting in increased activation of glioma-associated oncogene homolog (GLI) transcription factors. Uncontrolled disinhibition of SMO can result in failure of GCPs to differentiate, resulting in continued and excessive GCP proliferation, thereby overpopulating the IGL (Goodrich et al., 1997).

Mutations of genes encoding SHH pathway proteins are implicated in several human neurodevelopmental disorders (Hahn et al., 1996). Holoprosencephaly (HPE), most frequently caused by mutations of SHH, are the most common congenital forebrain abnormality in humans (Nanni et al., 1999; Solomon et al., 2010; Weiss et al., 2018a). Less commonly, mutations of PTCH1, the receptor for SHH, are also associated with HPE (Ming et al., 2002). Patients with HPE present with a wide array of phenotypes including forebrain malformations, craniofacial defects, and behavioral abnormalities including attention deficit hyperactivity disorder (ADHD) (Croen et al., 1996; Heussler et al., 2002).

Mutations of PTCH1 causing dysregulation of SHH are also associated with basal cell nevus syndrome (BCNS) (Gloude et al., 2016; Okamoto et al., 2014). Some BCNS patients develop skeletal anomalies, and approximately 3% develop the cerebellar brain cancer medulloblastoma (MB) (Lacombe et al., 1990). Molecular characterization of MB tumors has revealed that that 20-30% of MB patients have mutations in the SHH pathway (Northcott et al., 2012). The role of dysregulated SHH signaling in MB has been experimentally demonstrated using genetically modified strains of mice containing constitutively active Smo alleles, or knockout mutations of Ptch, that result in aberrant GLI1 transcriptional activity that gives rise to MB (Dey et al., 2012; Goodrich et al., 1997; Grammel et al., 2012; Hallahan et al., 2004; Lee et al., 2007; Schüller et al., 2008; Uziel, 2005; Wetmore et al., 2001). Ptch1-knockout mice, initially developed to evaluate the role of the SHH-PTCH signaling pathway in BCNS, have been used widely to study MB (Goodrich et al., 1997; Nitzki et al., 2012). Whereas homozygous knockout of the PTCH1 receptor gene (*Ptch1^-/-^*) is embryonically lethal due to failures of neural tube closure, haploinsufficient *Ptch* heterozygous mice (*Ptch1^+/-^*) are viable but have increased SMO and GLI1 activity resulting from decreased PTCH1 protein expression (Goodrich et al., 1997). Phenotypically, that dysregulation of GLI1 transcriptional programming results in increased GCP proliferation, aberrant neuronal migration, and MB in 10-20% of *Ptch1^+/-^* adult mice (Zurawel et al., 2000). In addition to cerebellar dysregulation and MB tumorigenesis, altered hippocampal structures have been reported in male *Ptch1^+/-^* adult mice (Antonelli et al., 2018). A finding consistent with SHH signaling also having potential roles in neurogenesis of hippocampal progenitors hippocampal (Yao et al., 2016).

As was observed in experimental mouse models, mutations in humans causing unregulated activation of SHH-signaling, either via gain of SMO or loss of PTCH function, also result in MB. Medulloblastoma are one of the most common solid tumors of childhood, accounting for approximately 20% of all pediatric tumors. There are noted sex differences in some molecular subgroups of MB, with overall incidence showing a 1.5:1 male to female sex ratio; however, the sex ratio differs by population and depends on the etiology of the tumor (Northcott et al., 2012; Sun et al., 2015). The WNT and SHH subgroups of MB present with a 1:1 sex ratio (Northcott et al., 2012). Estrogen receptor β (ERβ) is expressed in maturing GCPs and MB tumor cells and can modify MB growth and progression (Belcher, 2008; Jakab et al., 2001). Increased estrogen receptor signaling during cerebellar development and MB progression results in upregulation of cytoprotective ERβ-dependent insulin-like growth factor signaling that impacts GCP maturation and migration, and can increase MB tumor growth rate (Belcher, 2008; Cookman and Belcher, 2015).

Clinical treatment for MB is extremely aggressive and associated with severe life-long side effects in survivors. Typical treatment for MB involves primary tumor resection followed by radiation therapy and cytotoxic chemotherapy. Neurological complications including impaired attention and processing speed, learning and memory, language, visual perception, and executive functions occur in nearly all MB survivors and cause difficulties with social functions that can greatly decrease quality of life (Ribi et al., 2005). While associated with treatment, these behavioral defects and cognitive deficits resemble the hallmark behavioral symptoms associated with HPE and BCNS. The neurological deficits in MB survivors have been solely attributed to therapeutic side effects, but some component of the behavioral deficits in patients with MB may result from IGL overgrowth inherent to the SHH MB sequalae. To determine whether heterozygous mutation of *Ptch1* alone may influence behavior, male and female *Ptch1^+/-^* mice and their wildtype littermates were assayed using several behavioral tasks. Following behavioral analysis, brains from these *Ptch1^+/-^* mice were examined histologically to identify neuroanatomical alterations in structures potentially related to cerebello-cortical circuitry that could influence social behavior.

## 2. Experimental Procedures

### 2.1. Animal Husbandry

All animal procedures and reporting adhere to the ARRIVE guidelines (Supplemental data) and were performed in accordance with protocols approved by the North Carolina State University (NCSU) Institutional Animal Care and Use Committee following recommendations of the Panel on Euthanasia of the American Veterinary Medical Association. Study animals were housed on a 12:12 light cycle at 25°C and 45%-60% average relative humidity in an AAALAC accredited animal facility. Mice were housed in thoroughly washed polysulfone cages with woodchip bedding and pulped virgin cotton fiber nestlets (Ancare, Bellmore, NY). Soy-free Teklad 2020X diet (Envigo, Madison, WI) was supplied *ad libitum*. Sterile drinking water produced from a reverse osmosis water purification system (Millipore Rios with ELIX UV/Progard 2, Billerica, MA) was supplied *ad libitum* from glass water bottles with rubber stoppers and metal sippers.

Strains C57Bl6/J and STOCK *Ptch1^tm1Mps^*/J were obtained from Jackson Laboratory (Bar Harbor, ME). Breeding was performed in-house to propagate both lines. Heterozygote *Ptch1* mutants and wildtype littermates were used for experimental procedures. Adult mice that presented with MB tumors were excluded from analyses. Genomic DNA was isolated from a 5 mm tail biopsy using a rapid digestion method where 95 μl of lysis buffer reagent (Viagen Biotech, Los Angeles, CA; Cat: 102-T) and 5 μl of Proteinase K (20 mg/ml; Viagen Biotech, Los Angeles, CA; Cat: 501-PK) were added to the biopsy sample, incubated for 4 hours at 55°C and 45 minutes at 85°C. Isolated DNA was used to identify offspring genotype following recommended genotyping protocol (Jax, Bar Harbor, ME) and analyzed by agarose gel electrophoresis on 1.5% gels. Heterozygotes were identified by the presence of both wild-type (200 base pair) and mutant (479 base pair) specific PCR products. Mice were weaned on postnatal day 21 (PND 21), assigned a coded identification number (Supplemental Table 1), and identified by ear notching. No notable pathology differences or morbidity were detected in *Ptch1^+/-^* mice aside from development of MB in a subset of animals (18% of female; 9% of male) that were excluded from analyses due to death or ataxia associated with MB-like tumors detected at necropsy.

### 2.2. Behavior Tests and Analysis

All behavioral testing was conducted in a dedicated behavioral testing room at the NCSU Biological Resources Facility. Mice were transported in their home cage to the testing room on a covered rolling cart and tested after a minimum 30-minute acclimation period. Animals were tested in the last three hours of the light cycle. Both novel social and partner preference tasks used a blue, opaque arena (58 cm x 58 cm) with four high walls (43 cm). All arenas were in the same room with two conspecific animals run concurrently. Each animal was gently placed in the middle of the arena and given 30 minutes to explore, during which time they were not disturbed. No observer was present in the room. Behavioral data was digitally recorded (Handycam HDR-CX190, Sony, Tokyo) and automatically scored using TopScan behavioral analysis software (CleverSys, Inc, Reston, VA). For analysis, the floor of the open field arena was digitally divided into a 3×3 square grid, creating 9 squares in total of equal size. The middle square was designated as the center. All data was collected and analyzed by observers blinded to experimental group, and were validated by hand-scoring using a stop-watch.

#### 2.2.1. Novel Social Task

Adult mice (wild-type: mean age: PND96, range: PND75-114; *Ptch1^+/-^* mean age: PND99; range: PND75-119) were tested for 30 minutes in the novel social arena as described previously (Winslow, 2003). A naïve same sex and younger/smaller, unrelated wild-type stimulus animal was caged in a randomly selected corner of the arena. The first five minutes of the trial were excluded from analyses to account for increased exploratory behavior that is commonly observed during entry into a novel environment (Bailey and Crawley, 2009). Maximum velocity and distance traveled were used to examine motor function decrements. Center crosses, time spent along the wall, number of interactions, and time spent interacting were used to assess sociability. Latency to enter the stimulus animal’s area, frequency of movement within areas of the arena, number of contact bouts, and duration within stimulus animal’s area and exploratory areas were analyzed.

#### 2.2.2. Partner Preference Task

Twenty-four hours after the novel social testing, animals were tested with two same-sex conspecific wild-type animals to examine the effects of *Ptch1* haploinsufficiency on social interaction and formation of partner preference (Winslow, 2003). Briefly, the same stimulus animal used for the novel social task was caged in one corner as a now-familiar animal, while a novel unfamiliar mouse was caged in the opposite corner. Latency to enter each stimulus animal’s area, frequency of movement within areas of the arena, number of contact bouts with each animal, and duration within stimulus animal’s area and exploratory areas were analyzed.

#### 2.2.3 *Olfactory* Task

The ability of a subset of experimental animals to respond to a desirable olfactory cue was evaluated using an established protocol to assess intact recognition of a food-reward smell (Yang and Crawley, 2009). Briefly, animals were fasted for 12 hours overnight with *ad libitum* access to water and then placed into a clean cage containing a single previously buried Apple Jack (Kellogg, Battle Creek); the time required to locate and pick up the Apple Jack was measured using a stopwatch. An a priori maximum latency threshold of >900 seconds for food-treat discovery was defined as indicative of a decrement responsiveness (Yang and Crawley, 2009).

### 2.3. Neuroanatomy and Histology

Following completion of behavioral testing mice were euthanized by CO_2_ asphyxiation. Brains were isolated by dissection, rapidly frozen on powdered dry ice, and stored at −80°C until prepared for analysis. Brains and cerebella were separately embedded in OCT (Fisher Scientific, Hampton, NH) and mounted directly onto cryostat chucks for cryosectioning. Serial mid-sagittal cryosections (20 μm) from the central vermis of each cerebellum and serial coronal cryosections (40 μm) of the brain were sectioned using a Leica Cryostat (Leica CM1900, Nussloch, Germany), mounted onto Superfrost plus slides (Fisher Scientific, Pittsburgh, PA), and stored at −80°C until histological processing.

For Nissl staining, sections were brought to room temperature and immersed in xylene (Fisher Scientific, Hampton, NH; Cat: X5-500) for 30 minutes, 100% ethanol (Fisher Scientific, Hampton, NH; Cat: 22-032-601) for 3 minutes, 95% ethanol for 3 minutes, and Milli-Q water for 2 minutes before staining with 0.2% Cresyl Violet (Fisher Scientific, Hampton, NH; Cat: AC405760025) for 12 minutes. Sections were then dehydrated by immersing in 95% ethanol for 30 seconds, 100% ethanol for 30 seconds, and 3 washes in Xylene for 1 minute each. Slides were then cover-slipped using Permount (Electron Microscopy Sciences, Hatfield, PA; Cat: 17986-01).

Stained sections were examined by an investigator blind to genotype and sex on a Nikon Eclipse 80i microscope using a DSFi1 CCD camera controlled with Digital Sight software (Nikon; Melville, NY). Digital bright field micrographs of sections from the most medial 100-microns of the vermis were collected using the 2x objective for analysis of cerebellar morphology. For IGL area quantification, the most medial 20-micron section was defined using consecutive serial sections and identified by using the following criteria: the 4^th^ ventricle protrudes towards lobule IX, deep cerebellar nuclei (fastigial nucleus, interposed nucleus, and dentate nucleus) are absent, and lobule X had a distinct nodulus. Coronal 40-micron cryosections of the cortex were assessed for gross morphometric differences at each of the following three landmarks. The first region was identified using the following criteria: ammon’s horn extends through the section, supramammilary and medial mammillary nuclei were present, and nucleus of Darkschewitsch was present (Bregma -2.88). The second region was identified using the following criteria: the 3^rd^ ventricle extends through section and arcuate nucleus was present; (Bregma -1.755). The third region was identified using the following criteria: CA3 appears circular, lateral ventricle is rhomboid with a tail (Bregma -1.06). Digital bright field micrographs were collected using 1x and 2x Nikon objectives (Nikon, Tokyo, Japan; Cat: MRL00012 and MRL00022) and compared to a standardized atlas to identify gross morphometric differences. (Allen Institute for Brain Science, 2011, Seattle, WA). Areas of interest were measured using Nikon NIS Elements AR 3.2 with final figures of representative sections generated using Adobe Photoshop (San Jose, CA).

### 2.4. Statistical Analysis

Data analysis was performed using a 2-way (sex, genotype) or 3-way (sex, genotype, novelty/sociality/5-min period) multivariate analysis of variance as indicated. For analysis of cerebellar lobules and cortical layers, data analysis was performed using repeated measure ANOVA. Litter was included as a covariate and adjusted for if necessary. All animal assignments and litter information are provided (Supplemental Table 1). If overall effects were significant, a Fisher’s least significant differences post hoc test was performed to evaluate pair-wise differences. For neuroanatomical endpoints, confounds related to possible litter effects were avoided by limiting analysis to one animal of a given sex from each litter. A minimal level of statistical significance for differences in values among or between groups was considered p < .05. Percentage data was arcsine transformed (arcsine of the square root of the percentage) prior to statistical analysis. All data were analyzed using GraphPad Prism v8 (GraphPad; La Jolla, CA) or SPSS v26 (IBM, Armonk, NY).

## 3. Results

### 3.1. General locomotor function and exploratory behavior

Since the cerebellum is critical in motor function, general locomotor function was assessed during the open-field novel social task to evaluate whether *Ptch1^+/-^* mice exhibit motor deficits. Multivariate analysis of variance (MANOVA) showed no deficits in maximum velocity over the 30-minute trial (F(1, 68) = .14, p = .71) (Figure 1A). Multivariate analysis of variance showed an interaction of sex and genotype on total distance traveled (F(1, 54 = 4.52, p = .04, η2 = .55)). Post hoc analysis using Fisher’s LSD indicated that total distance traveled by *Ptch1^+/-^* females only was increased compared to wild-type (females: p = .009; males: p = .84) (Figure 1B). Repeated measures 3-way MANOVA with sex (male, female) as the within-subjects factor, 5-min period (1-6), and genotype (wild-type, *Ptch1^+/-^*) as the between-subjects factors revealed a significant interaction of sex and genotype on distance traveled across 5-minute periods (F(1, 53 = 4.28, p = .04, η2 = .18)). A follow-up repeated measures 2-way MANOVA within sex revealed that females only were significantly different in distance traveled across 5-minute periods (females: p = .002; male: p = .47) (Figure 1C and 1D).

**Figure 1.**
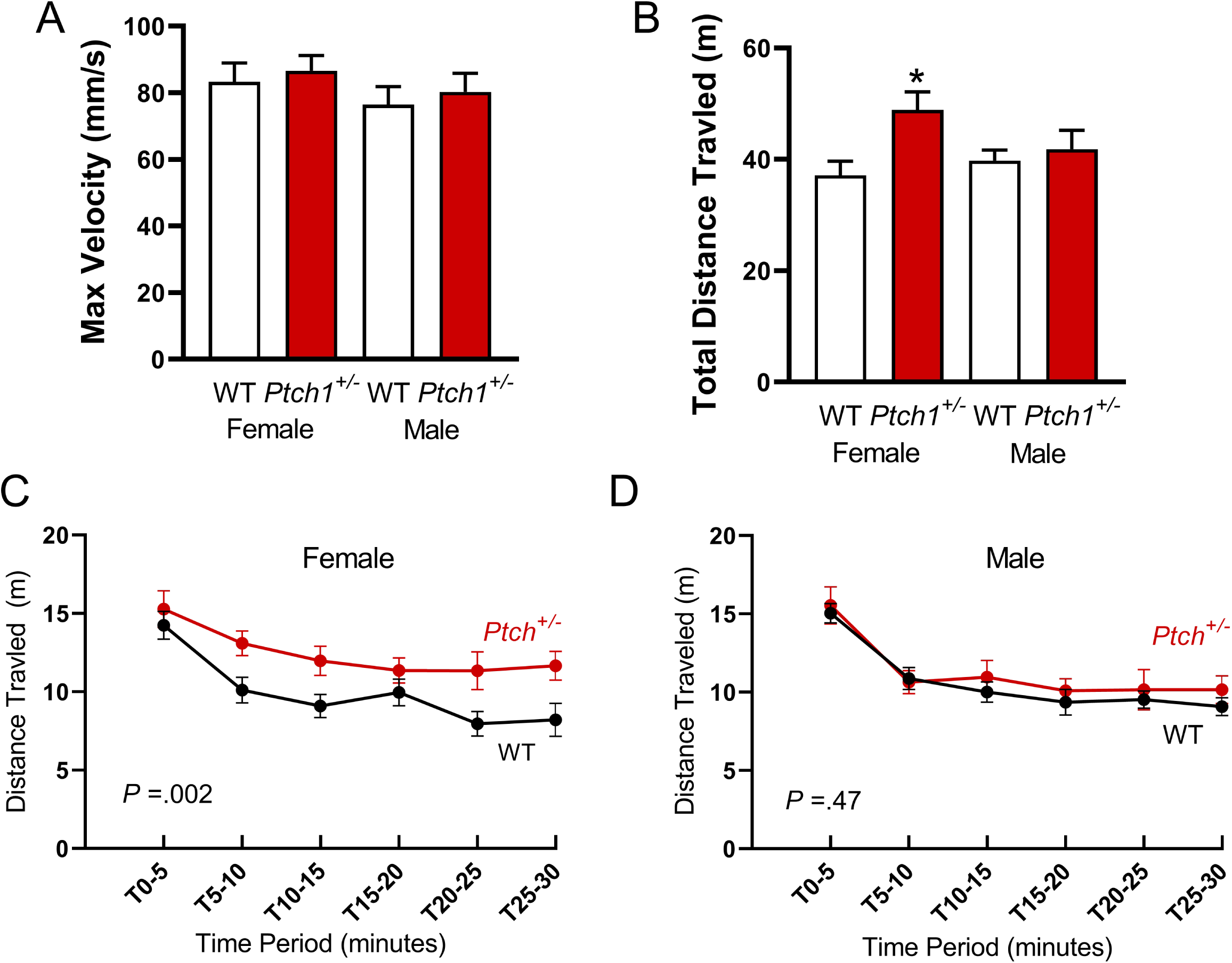
Behavioral assessment using a novel open-field task. Maximum velocity (A), total distance traveled (B), and distance traveled over 5-min periods in females (C) and males (D) are shown. Values are expressed as mean ± SEM. * = p < .05. Final animal numbers animals used for novel social task follow. Females: wild-type, n=18; *Ptch1^+/-^*, n=18. Males: wild-type, n=23; *Ptch1^+/-^*, n=13. Three wild-type females were excluded because they climbed atop the holding cup and avoided the task.

During the open-field novel social task, time spent along the wall and number of center crosses over the 30-minute trial were assessed to examine exploratory behavior. Multivariate analysis of variance showed no differences in time spent along the wall (F(1, 68 = .79, p = .38)) (Supplemental Figure 1). Consistent with increased distance traveled, MANOVA found a significant effect of genotype on center crosses (F(1, 68 = 8.47, p = .006, η2 = .81)) (Supplemental Figure 1). Post hoc analysis using Fisher’s LSD indicated a significant increase in number of center crosses in *Ptch1^+/-^* females only (females: p = .02; males: p = .55). In the novel social task, MANOVA found no differences in the number of interactions (F(1, 68 = 1.98, p = .16) or the duration of those interactions (F(1, 68 = 0.20, p = .66) with the novel animal.

### 3.2. Social behavior

To assess the effects of *Ptch1* mutation on social behavior, a partner preference task was used. As in the novel social task, MANOVA of the partner preference task showed no differences in maximum velocity (F(1, 57 = .59, p = .45) (Figure 1A) nor time spent along the wall (F(1, 57 = 3.72, p = .06)) (Supplemental Figure 2). In the partner preference task, MANOVA again detected a significant interaction between sex and genotype on total distance traveled (F(1, 56 = 4.12, p = .048, η2 = .51). Post hoc analysis using Fisher’s LSD identified a significant increase in total distance traveled was again in *Ptch1^+/-^* females only (females: p = .002; males: p = .71). Multivariate analysis of variance again showed a main effect of genotype on center crosses consistent with increased distance traveled (F(1, 57 = 5.99, p = .02, η2 = .67)) (Supplemental Figure 2). Post hoc analysis using Fisher’s LSD identified a significant increase in total distance traveled was again in *Ptch1^+/-^* females only (females: p = .02; males: p = .22).

A 3-way MANOVA with sex (male, female) as the within-subjects factor, and novelty (novel, familiar) and genotype (wild-type, *Ptch1^+/-^*) as the between-subjects factors, revealed an interaction between sex and genotype on the duration of interactions during the partner preference task (F(1, 110) = 8.59, p = .005, η2 = .82)). Post hoc analysis using Fisher’s LSD revealed that *Ptch1^+/-^* females only show increased time spent interacting with both novel (females: p = .047; males: p = .72) and familiar animals (females: p = .03; males: p = .93) (Figure 2A). All other main effects were non-significant and not relevant to the tested hypotheses.

**Figure 2.**
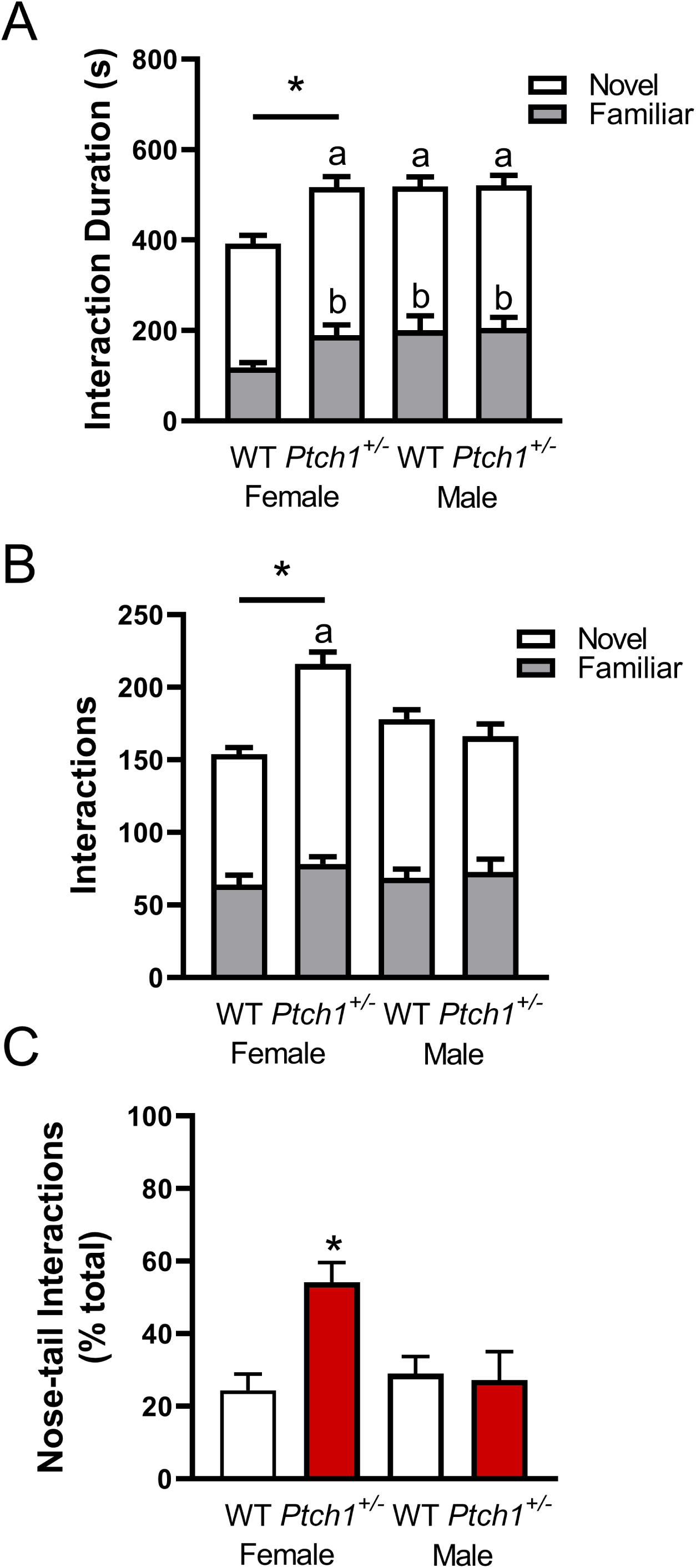
Behavioral assessment using a partner preference task. Duration of interactions with novel and familiar animals (A), the number of those interactions (B), and the type of those interactions (C) are shown. Lower-cased (a) and (b) illustrate significant differences within each level of the ANOVA compared to wild-type females. Values are expressed as mean ± SEM. * = p < .05. Final animal numbers animals used for partner preference task follow. Females: wild-type, n=17; *Ptch1^+/-^*, n=14. Males: wild-type, n=19; *Ptch1^+/-^*, n=12. One wild-type female that climbed atop the holding cup and avoided performing the task was excluded.

A 3-way MANOVA with sex (male, female) as the within-subjects factor and novelty (novel, familiar) and genotype (wild-type, *Ptch1^+/-^*) as the between-subjects factors revealed an interaction between sex and genotype on the number of interactions during partner preference task (F(1, 111) = 14.52, p = .0002, η2 = .22)). That interaction was qualified by an interaction between novelty, sex, and genotype (F(1, 111) = 7.698, p = .007, η2 = .11)). Post hoc analysis using Fisher’s LSD revealed that *Ptch1^+/-^* females only show an increase in the number of interactions with the novel animal (females: p < .0001; males: p = .10) (Figure 2B). All other main effects were non-significant and not relevant to the tested hypotheses.

A 3-way ANOVA with sex (male, female) as the within-subjects factor, and sociality (nose-nose, nose-tail) and genotype (wild-type, *Ptch1^+/-^*) as the between-subjects factors, revealed a significant interaction between sex, sociality, and genotype on the percentage of nose-tail bouts (F(1, 136) = 7.24, p = .01, η2 = .75). Post hoc analysis using Fisher’s LSD revealed that *Ptch1^+/-^* females only show increased nose-tail bouts relative to wild-type (females: p = .0003; males: p = .81) (Figure 2C). All other main effects were non-significant and not relevant to the tested hypotheses.

### 3.3. Olfaction

The ability of test animals to respond to a desirable food-treat was evaluated to ensure that modifications in social behaviors were not influenced by unanticipated decrements in ability to identify and respond to olfactory cues or a lack of motivation. A subset of wild-type (female: n=7; male: n=5) and *Ptch1^+/-^* (female: n=3; male: n=2) were evaluated using an established protocol to assess intact recognition of smell wherein a latency greater than 15 minutes (900 seconds) is considered indicative of a decrement in smell (Yang and Crawley, 2009). There were no test failures with all animals tested rapidly discovering the olfactory stimulus in less than the allotted task time. The maximum time for any animal to complete the task was less than 4 minutes (231 seconds).

### 3.4. Cerebellar and cortical structures

In contrast to the stereotypic morphology of the cerebellum observed in both male and female wild-type mice (Figure 3A and 3C), granule cell overgrowth with localized thickening of the IGL was commonly observed in the *Ptch1^+/-^* mutants (Figure 3B and 3D; asterisks). Ectopic granule cells were also observed in 43% (6/14) of *Ptch1^+/-^* females (Figure 3B; arrow) and 45% (5/11) of *Ptch1^+/-^* males, compared to vermis of the cerebellum from wild-type mice (Figure 3A, female; 3C, male) where ectopic granule cells were not observed in either sex (female: n=9; male: n=9).

**Figure 3.**
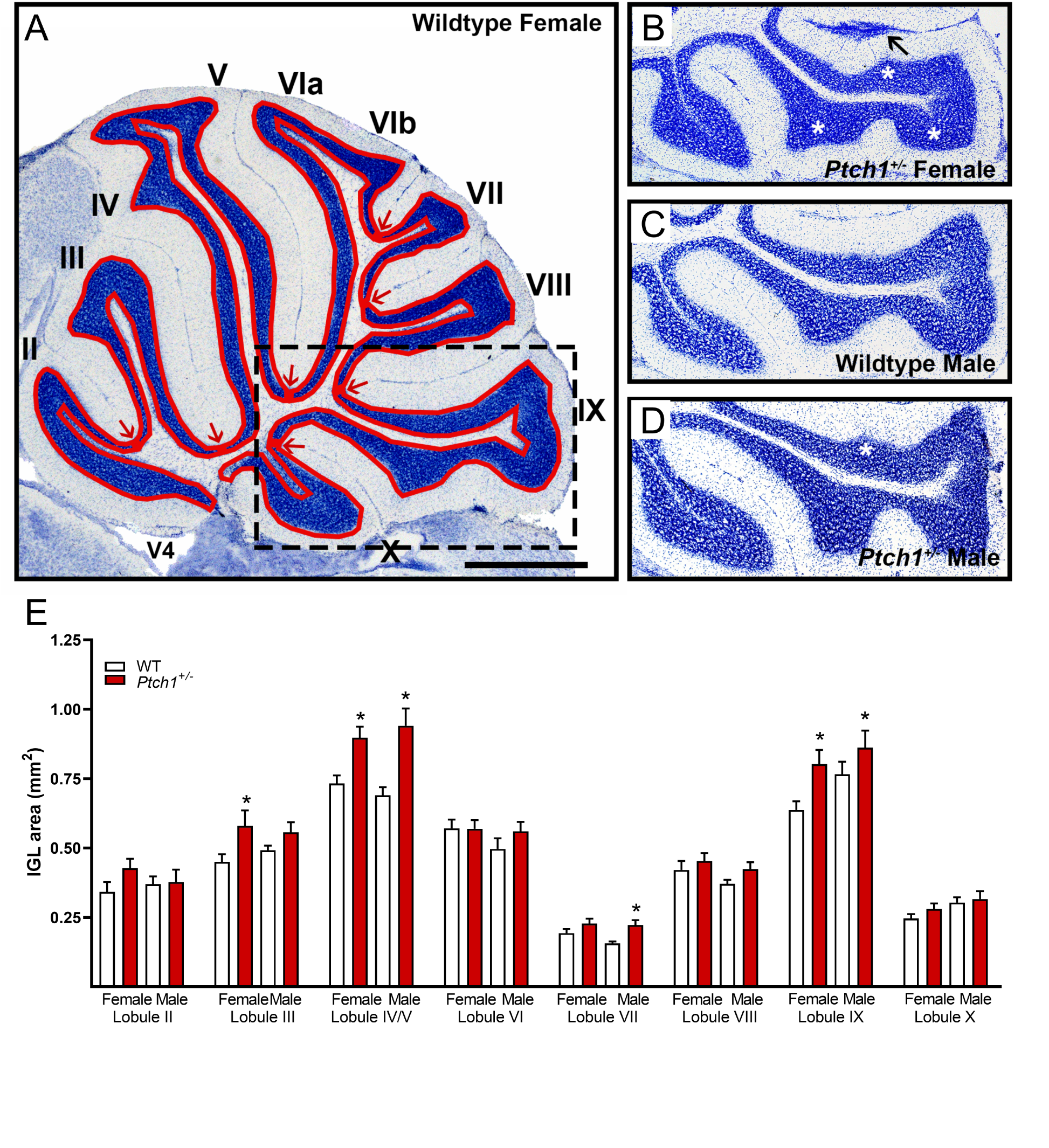
Assessment of changes in cerebellar structures. Sagittal cerebellar section (20 μm) showing lobular definition in a Wild-Type female, with cerebellar lobules (II – X) and the fourth ventricle labeled (V4). The lobular structures are outlined as quantified (A). The hatched outline in (A) denotes the region compared to a *Ptch1^+/-^* female (B), where differences in *Ptch1^+/-^* females are highlighted with an arrow indicating ectopic overgrowth and asterisks showing overgrowth of lobule IX (B). A wild-type male (C) and *Ptch1^+/-^* male (D) are also shown. Individual lobular IGL area are shown (E). Scale bar = 1 mm. Values are expressed as mean ± SEM. * = p < .05. Final animal numbers animals used for IGL area comparison follow. Females: wild-type, n=9; *Ptch1^+/-^*, n=5. Males: wild-type, n=9; *Ptch1^+/-^*, n=9.

The extent of granule cell overgrowth was assessed by quantitative comparison of IGL area in the cerebellar vermis of wildtype and *Ptch1^+/-^* mutants. Repeated measures analysis of variance showed a significant effect of genotype on total IGL area (F(1, 31 = 7.349, p = .01, η2 = .99)). Post hoc analysis for total IGL area using Fisher’s LSD was not statistically significant in either sex (Figure 3B; female: p = .06; male: p = .07), but differences in the area of individual lobules of the cerebellum appeared to drive the detected effect of genotype (Table 1; Figure 3E). Repeated measures ANOVA for the IGL area of individual lobules detected a significant main effect of genotype on IGL area (F(1, 31 = 13.7, p = .002, η2 = .98) with post hoc analysis using Fisher’s LSD demonstrating specific effects in both sexes on lobules IV/V (females: p = .02; males: p = .001), VII (females: p = .04; males: p = .009), VIII (females: p = .046; males: p = .008), and IX (females: p = .001; males: p = .01) and in males only in lobule VI (females: p = .82; males: p = .001) (Figure 3E). Overt loss of Purkinje cells or differences in the width of the Purkinje cell monolayer were not observed in the cerebellum of either sex.

**Table 1.** Internal Granule Cell Layer Area

| Lobule | Wild-Type | *Ptch1*<sup>+/-</sup> | Wild-Type | *Ptch1*<sup>+/-</sup> |
| --- | --- | --- | --- | --- |
|  | Female |  | Male |  |
| II | 0.342 ± 0.035 | 0.427 ± 0.033 | 0.369 ± 0.028 | 0.377 ± 0.044 |
| III | 0.450 ± 0.027 | 0.580 ± 0.055 | 0.492 ± 0.017 | 0.556 ± 0.037 |
| IV/V | 0.732 ± 0.029 | **0.897 ± 0.040\*** | 0.690 ± 0.029 | **0.941 ± 0.062\*** |
| VI | 0.570 ± 0.031 | 0.569 ± 0.031 | 0.468 ± 0.034 | **0.616 ± 0.046\*** |
| VII | 0.193 ± 0.015 | **0.227 ± 0.018\*** | 0.156 ± 0.007 | **0.222 ± 0.017\*** |
| VIII | 0.421 ± 0.032 | **0.453 ± 0.028\*** | 0.371 ± 0.015 | **0.424 ± 0.024\*** |
| IX | 0.637 ± 0.031 | **0.803 ± 0.051\*** | 0.765 ± 0.045 | **0.861 ± 0.061\*** |
| X | 0.246 ± 0.015 | 0.280 ± 0.019 | 0.303 ± 0.019 | 0.316 ± 0.028 |

Measurements expressed as mean ± SEM (mm<sup>2</sup>). \* = p < .05.

Outside of the cerebellum, malignancy or gross morphological changes in the brains of the *Ptch^+/-^* mutants were not observed. To evaluate potential structural differences in the extra-cerebellar brain regions of *Ptch1^+/-^* mutants, hippocampal and cortical structures implicated in playing a functional role in cerebellar functions and related behaviors were compared. Analysis of variance of the length of ammon’s horn (Figure 4A-4C) showed a statistically significant interaction of genotype and sex (F(1, 14 = 4.62, p = .0495, η2 = .39). Post hoc analysis using Fisher’s LSD indicated that this hippocampal structure in *Ptch1^+/-^* females only was significantly smaller than wildtype (females: p = .006; males: p = .75) (Figure 4D). Analysis of variance of the dentate gyrus length showed a statistically significant interaction of genotype and sex (F(1, 14 = 7.16, p = .02, η2 = .40)). Post hoc analysis using Fisher’s LSD indicated that the dentate gyrus of *Ptch1^+/-^* females only was similarly reduced in size compared to wild-type (females: p = .006; males: p = .71) (Figure 4D). Lateral ventricle area was measured because of noted differences of the lateral ventricle in BCNS syndrome and ASD (Shiohama et al., 2017; Turner et al., 2016). Analysis of variance showed no differences in lateral ventricle area (F(1, 11 = 0.93, p = .35)).

**Figure 4.**
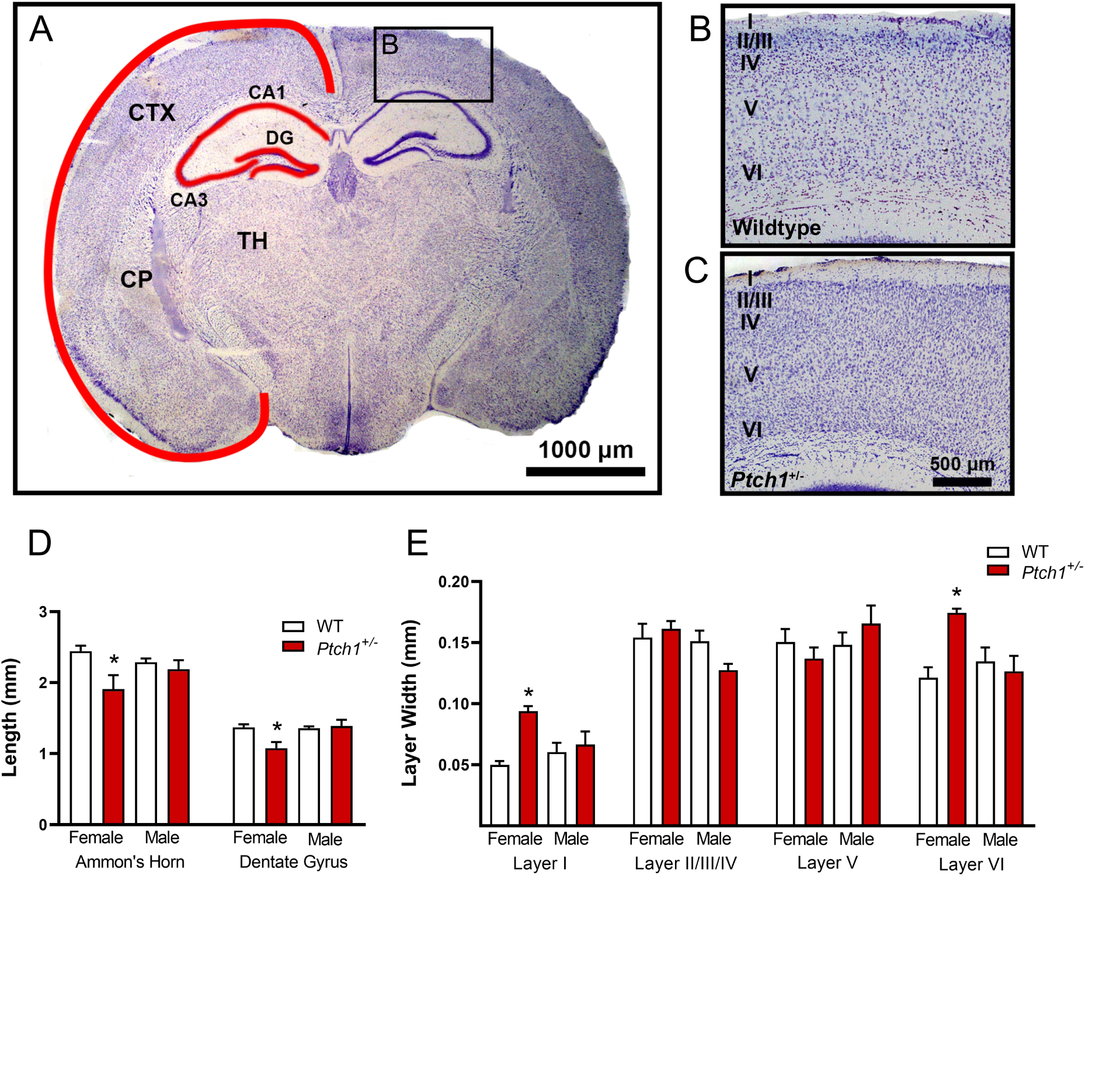
Assessment of changes in cortical structures. Coronal brain section (40 μm) highlighting measured regions of cortex (CTX), ammon’s horn (CA1, CA3), dentate gyrus (DG) (A) with region identified using caudate putamen (CP) and thalamus (TH) as landmarks. Inset image shows cortical layers (labeled I, II-IV, V, and VI of WT female (B) and *Ptch^+/-^* female mouse (C). Ammon’s horn (CA1-3), dentate gyrus, and cerebral cortex are outlined in red. Cortical layers are labeled I, II-IV, V, and VI. Scale bar = 1 mm (A) and 500 μm (B,C). Expressed as mean ± SEM. * = p < .05. For overall cortical layer length and width, the number of analyzed animals follow. Females: wild-type, n=4; *Ptch1^+/-^*, n=4. Males: wild-type, n=6; *Ptch1^+/-^*, n=4. Final animal numbers for individual cortical layers follow. Females: wild-type, n=4; *Ptch1^+/-^*, n=3. Males: wild-type, n=6; *Ptch1^+/-^*, n=3.

In the cortex, a main effect of genotype on overall cortical layer length (F(1, 14 = 6.24, p = .03, η2 = .80)) was detected, however post hoc analysis of total cortical layer length (Figure 4A) was not statistically significant for either sex (female: p = .10; male: p = .10). Analysis of variance of overall cortical layer width showed no effect from sex or genotype (F(1, 14 = 1.65, p = .22). However, increased cellularity in all cortical layers was observed in *Ptch1^+/-^* females (Figures 4B; 4C). Repeated measures ANOVA found a significant effect of genotype on the width of individual cortical layers (F(1, 25) = 8.2, p = .02, η2 = .48). Post hoc analysis using Fisher’s LSD detected specific effects in lobule VI in females only (females: p = .007; males: p = .35) (Figure 4E). No effect of sex or genotype was detected in either sex in cortical layers I (females: p = .22; males: p = .65)), II/III/IV (females: p = .09; males: p = .70)) or V (females: p =.14; males: p = .20).

## 4. Discussion

### 4.1. Sex-specific alterations of social behavior and activity of Ptch1^+/-^ females

Dysregulation of SHH signaling resulted in a significant interaction of sex and genotype that drove the female-specific social behavioral effects observed in the novel open-field and partner preference tasks. *Ptch1^+/-^* females traveled further and had more center crosses in both the novel open-field and partner preference tasks. In the partner preference task, *Ptch1^+/-^* females spent increased time with both novel and familiar animals and had increased nose-tail interactions relative to wild-type mice. In mice, most novel social interactions involve nose-nose contact, this investigative social activity contrasts with aggressive nose-tail interactions (Silverman et al., 2010). There were no detected differences in the type or duration of social behavior displayed in the male *Ptch1^+/-^* mutants. Prosocial phenotypes observed in *Ptch1^+/-^* females indicated differential responsiveness to dysregulated SHH signaling wherein female sex exacerbates alterations in social behavior and hyperactivity.

Because the *Ptch* knockout mutation used in this study resulted in global deficits of PTCH, pleiotropic effects from *Ptch1* haploinsufficiency could have contributed to the observed behavioral changes. The cerebellum is critical in motor movement (Altman and Bayer, 1997), and disrupted SHH signaling has been linked to changes in olfactory neuron production (Daynac et al., 2016; Gomez et al., 2019; Ihrie et al., 2011; Tong et al., 2015). To rule out trivial explanations for altered behaviors observed, we carefully assessed movement in our novel open-field task for evidence of decrements related to compromises mechano-skeletal ability, cerebellar control of coordination or movement, and to rule out defects related to olfaction. No deficits in movement velocity, distance traveled, or latency to find an olfaction cue were detected. In fact, there was a paradoxical increase in the activity of *Ptch1^+/-^* females indicated by increases in total distance moved. Sustained activity after the first five minutes indicates a failure of *Ptch1^+/-^* females to acclimate to the arena (Figure 1C), suggesting more complicated sex-specific impacts than decreased movement coordination or loss of the ability to smell were responsible for the observed alterations in behavior.

Increased activity and prosocial behaviors of *Ptch1^+/-^* females could be explained by a hyperactivity phenotype. In humans, ADHD has been associated with HPE caused by SHH mutations primarily in female humans with noted high intellectual function (Heussler et al., 2002; Solomon et al., 2012), demonstrating that altered SHH signaling can lead to hyperactivity. Similar to the sex differences observed in *Ptch1^+/-^* female mice, human females are also at increased risk for diagnosis of HPE with sex ratios ranging from 1.2:1 to 2.3:1 female:male (Croen et al., 1996; Mouden et al., 2016; Weiss et al., 2018a). The prosocial phenotype observed in *Ptch1^+/-^* females may be due to a mild sex-specific HPE-like hyperactive phenotype resulting in indiscriminate social interactions.

### 4.2. Neuroanatomical examination of cerebellar structure

The presented findings from behavioral tasks supplement previous work demonstrating that haploinsufficiency of Ptch1 can alter cognitive behaviors including reduced motor learning ability and performance on a spatial memory task by linking dysregulation of SHH signaling with hyperactivity and altered social behaviors (Antonelli et al., 2018; Dutka et al., 2015). The altered social behavior in *Ptch1^+/-^* females may be related to cerebellar overgrowth, which is increasingly appreciated for its role in schizophrenia, dementia, and other psychiatric disorders (Phillips et al., 2015).

Imaging, clinical, and experimental animal studies have linked the cerebellum with cognitive brain disorders including ASD and BCNS syndrome (Becker and Stoodley, 2013; Lo Muzio, 2008), with cerebellar hypoplasia and reduced Purkinje cell numbers most commonly linked with ASD (Bauman, 1991; Palmen et al., 2004). However, investigations mechanistically linking the cerebellum to cognition and circuits modulating social behavior are preliminary and deserve further attention (McKimm et al., 2014). There were no detectable differences in the width of the Purkinje cell monolayer in *Ptch1^+/-^* mice or overt losses of Purkinje cells. Increases in IGL area was generally evident across the entire cerebellum in both male and female *Ptch1* heterozygotes, indicating that cerebellar phenotypes are likely related to increased proliferation and density of granule cells that resulted in detectable increases in area of some cerebellar lobules. While IGL area was generally increased across all lobules, significant cerebellar overgrowth in the IGL of lobules IV/V and IX was found in both sexes, whereas GCP overgrowth of lobule III reached significance in females and overgrowth of lobule VII was significant in males. Lobules IV/V and IX of the cerebellum have previously been identified as sensitive to SHH pathway defects via mutation of Gli1, Gli2, and Smo (Corrales et al., 2006; Tan et al., 2018); both *Ptch1^+/-^* males and females developed GCP overgrowth specifically in these two lobules.

The cerebellum is divided into functionally specific regions with lobules I-V and IX often associated with motor function and working memory in rodents and humans (D’Mello et al., 2015; Guell et al.; Lawrenson et al., 2018). Reductions in volume of lobules IV/V and IX are also associated with social impairments in patients with autism (D’Mello et al., 2015). Vermal hyperplasia of cerebellar lobules VI and VII in a subgroup of autism patients (Courchesne et al., 1994), and Purkinje cell hyperplasia in lobules IV/V and IX associated with impaired social behavior have been described (Cupolillo et al., 2016). Because both hypoplasia and overgrowth of the different layers of the cerebellum are linked to altered social behavior, an appropriate balance of Purkinje and granule cell inputs are likely necessary for normal cerebellar modulation of behavior; vermal granule cell hyperplasia in *Ptch1^+/-^* females may be driving the observed alterations in social behavior and hyperactivity-like phenotype.

Treatment for SHH MB is extremely aggressive and involves primary tumor resection followed by high-dose chemotherapeutics (Kumar et al., 2017). Following treatment, behavioral decrements of attention and processing speed, learning and memory, language, visual perception, and executive function are common (Ribi et al., 2005). These neurological complications, attributed entirely to therapeutic side effects, occur in nearly all MB survivors and resemble behavioral defects and cognitive deficits observed in cerebellar-associated disorders including HPE, BCNS, and William’s Syndrome (Lo Muzio, 2008; Reiss et al., 2004; Weiss et al., 2018b). Considering that *Ptch1^+/-^* females exhibit altered behavior, some neurological complications noted in MB survivors may exist prior to treatment as disease etiology rather than solely a consequence of adverse effects of treatment, and those neurological symptoms could be different in males and females. Understanding of the behavioral sequelae inherent to, and the innate sex differences within, MB etiology is critical to improving the treatment plan for MB patients.

### 4.3. Cerebellar modulation of social behavior

Consistent with the association between reduced hippocampal size, hyperactivity, learning, and memory deficits in humans (Al-Amin et al., 2018; Tamnes et al., 2014), *Ptch1^+/-^* female mice exhibited changes in hippocampal structures associated with altered behavior. Altered hippocampal structures have previously been reported in *Ptch1^+/-^* male mice, although females were not examined (Antonelli et al., 2018). SHH signaling directs neurogenesis and is a critical player in hippocampal plasticity (Yao et al., 2016). *Ptch1^+/-^* females also exhibited increased cellularity and thickness of cortical layer VI.

Cortical layer VI is the earliest developing cortical layer (Gilmore and Herrup, 1997) and has the greatest diversity of neuronal cell types, with many excitatory pyramidal neurons, glutamatergic neurons, spiny stellate neurons that project to the thalamus, and local inhibitory neurons (Briggs, 2010). SHH signaling is critical to mitogenesis and speciation of cortical progenitors early in development (Yabut and Pleasure, 2018) and SHH signaling induces cortical growth (Wang et al., 2016). The observed decrement in hippocampal size and increased cortical layer thickness resulting from decreased SHH signaling provides further evidence that neuronal changes outside the cerebellum may contribute to noted behavioral changes in *Ptch1^+/-^* females. The observed behavioral changes in activity and social behavior could be related to decreases in hippocampal plasticity and the alteration of projections to extracerebellar circuits. Alternatively, the observed changes in cortical and hippocampal structures may result from an imbalance of extracerebellar inputs or activity to these structures resulting from the indirect influence of increased cerebellar granule cell numbers due to increased synaptic inputs on Purkinje cells and subsequent impacts on extracerebellar activity.

A potential mechanism linking abnormal cerebellar pathology to impaired social function is via cerebellar modulation of dopamine release within the mPFC (Rogers et al., 2013b). The mPFC incorporates cerebellar output through the cerebellar dentate nucleus which modulate dopamine release in the mPFC via two primary pathways that appear to contribute equally to mPFC activation: 1) contralateral glutamatergic projections from the cerebellar dentate nucleus to reticulotegmental nuclei that project to pedunculopontine nuclei and stimulate mesocortical dopaminergic neurons in the ventral tegmental area (VTA), which then project to the mPFC, or 2) contralateral glutamatergic projections of the cerebellar dentate nucleus project to thalamic mediodorsal and ventrolateral nuclei (ThN md and ThN vl) that send glutamatergic effects to the mPFC to modulate mesocortical dopaminergic terminal release in the mPFC via excitatory glutamatergic synapses (McKimm et al., 2014) (Figure 5).

**Figure 5.**
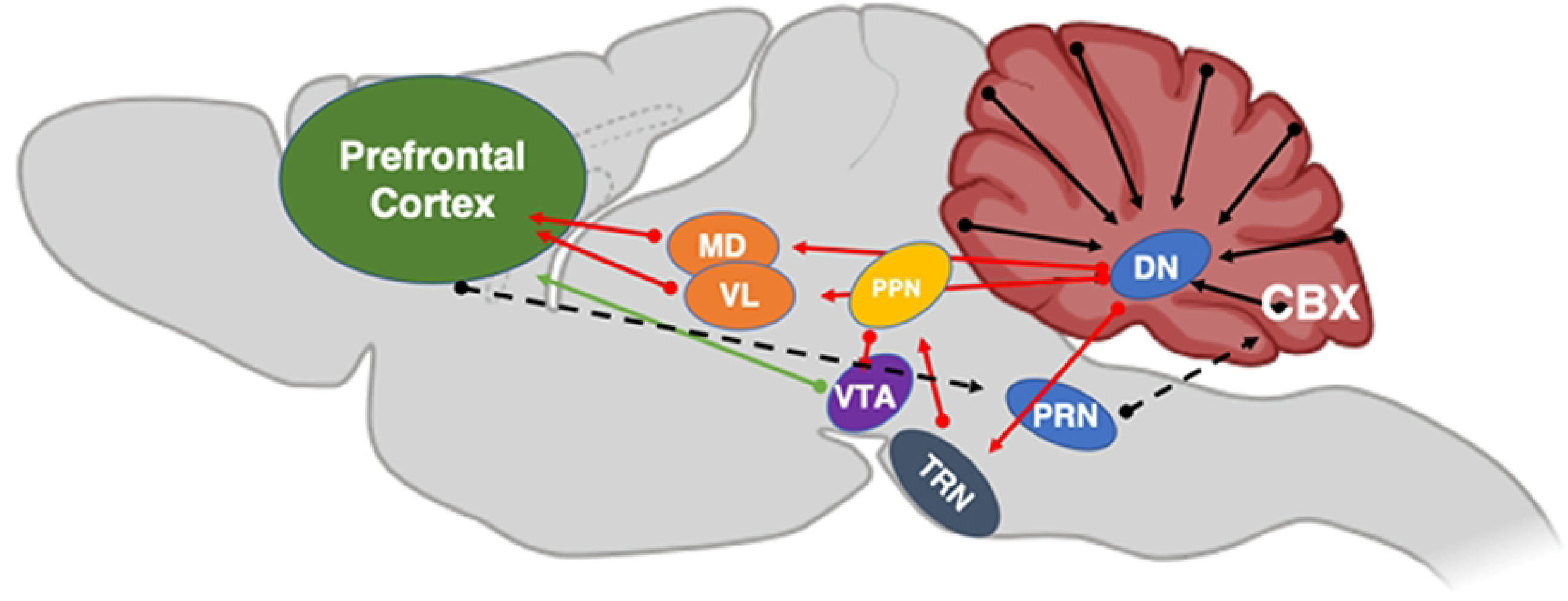
Proposed pathway of cerebello-cortical circuitry. Shown is a cartoon of the proposed neuronal circuits linking outputs from the cerebellar cortex (CBX) to projections from deep cerebellar nuclei (DN) to neurons in the prefrontal cortex (PFC) via the tegmental reticular nuclei (TRN), pedunculopontine nuclei (PPN), the ventral tegmental area (VTA), thalamic mediodorsal (MD) and ventrolateral (VL) nuclei, and pontine reticular nuclei (PRN).

Changes in projections through the VTA may explain why *Ptch1^+/-^* females display altered social behavior, because SHH is necessary for the specification of dopamine cell fate in the VTA (Blaess et al., 2011). Supporting this possibility, experimental mutation of the *Ptch* coreceptor, *Cdon* in mice, increased numbers of dopamine neurons in the VTA and eleveated levels of dopamine and its metabolites in the mPFC (Verwey et al., 2016). Thus, alterations in dopamine release within the mPFC might explain the increased activity and altered social behavior observed in female Ptch^+/-^ mice. Supporting this possibility, mice with conditional inactivation of Smo specifically in dopamine cells are hyperactive (Zhou et al., 2016), whereas dopamine receptor D1 blockade in the mPFC of rats reduces the distance traveled in an open-field task (Hall et al., 2009). Those studies suggest that cerebellar-related increases in mPFC dopamine release might mediate the alterations in activity and social observed in the *Ptch1^+/-^* females.

### 4.4. Female-specific effects of Ptch1 mutation

The more severe cerebellar overgrowth and behavior phenotypes observed in female *Ptch1^+/-^* mice could also be explained by increased responsiveness of GCPs to circulating estrogens. It is notable that in SHH mouse models of MB the incidence of tumors is higher in females (Svärd et al., 2009). Estrogens also play important roles in regulating cerebellar granule cell proliferation and MB progression (Guillette et al., 2018). Considering the progressive nature of MB development in *Ptch1^+/-^*, the apparent increased sensitivity of females to dysregulated SHH signaling could be related to the increased levels of circulating estrogens in mature females. These activities of estrogen are mediated through modulation of rapid estrogen signaling, estrogen receptor-regulated gene expression, and resulting modulation of growth factor-related signal transduction pathways (Cookman and Belcher, 2015; Garcia-Segura et al., 2006). Increased activation of ERβ signaling increases GCP mitogenesis and migration, and upregulates neuroprotective mechanisms in mature granule cells, which also act in the etiology and progression of MB to drive tumor progression (Belcher, 2008; Cookman and Belcher, 2015; Guillette et al., 2018). It is possible that the more sever phenotypes detected in *Ptch1^+/-^* female mice are related to a relative increase in estrogen signaling in ERβ positive *Ptch1^+/-^* granule cell-like precursors. It is also possible that sex-related differences in the proposed dopaminergic cerebellar-mPFC circuitry, leading to differential dopamine release or dopamine receptor activity in the mPFC, could sex-specifically contribute to differences in structural organization and behaviors that are associated with decreased SHH signaling.

## 5. Conclusion

The disruption of SHH-signaling in *Ptch1^+/-^* mice causes lobule-specific overgrowth in cerebellar structures previously linked to ASD in both sexes. Additionally, *Ptch1^+/-^* mice exhibited female-specific alterations in the hippocampus and cortical layers, hyperactivity, and altered social behaviors. Based on these findings, it is proposed that a subset of the behavioral phenotypes observed in MB patients following treatment are a component of MB sequalae rather than side effects of treatment. Further work focusing on the role of estrogen and SHH signaling cross-talk is needed to elucidate the developmental and functional nature of the proposed cerebellar-mPFC circuitry and the functional role of the cerebellum as a mediator of sex-specific behaviors.

## Acknowledgements

We are indebted to Dr. Heather Patisaul for assistance with the design and analysis of the behavioral studies, Dr. Theresa Guillette and the other members of the Belcher and Patisaul labs who assisted in various aspects of the study and gave critical feed-back on the manuscript. We would also like to express our great appreciation for the support of Sandy Elliott and the staff of the NCSU BRF who go above and beyond to facilitate our animal studies.

